# The Role of the Co-Chaperone DNAJB11 in Polycystic Kidney Disease: Molecular Mechanisms and Cellular Origin of Cyst Formation

**DOI:** 10.1101/2024.03.04.582938

**Authors:** Tilman Busch, Björn Neubauer, Lars Schmitt, Isabel Cascante, Luise Knoblich, Oliver Wegehaupt, Felix Schöler, Stefan Tholen, Alexis Hofherr, Christoph Schell, Oliver Schilling, Lukas Westermann, Anna Köttgen, Michael Köttgen

**Affiliations:** Department of Medicine IV - Nephrology and Primary Care, Faculty of Medicine and Medical Center, University of Freiburg, Freiburg, Germany; Institute for Immunodeficiency, Center for Chronic Immunodeficiency (CCI), Medical Center, Faculty of Medicine, University of Freiburg, Freiburg, Germany; Department of Pathology, Faculty of Medicine and Medical Center, University of Freiburg, Freiburg, Germany; Institute of Genetic Epidemiology, Faculty of Medicine and Medical Center - University of Freiburg, Freiburg, Germany; CIBSS - Centre for Integrative Biological Signalling Studies, University of Freiburg, Freiburg, Germany

**Author notes:** **Correspondence:** Anna Köttgen, MD, Institute of Genetic Epidemiology, Faculty of Medicine and Medical Center, University of Freiburg, Hugstetter Str. 49, 79106 Freiburg, Germany; Michael Köttgen, MD, Department of Medicine IV – Nephrology and Primary Care, Medical Center – University of Freiburg, Institute for Disease Modeling and Targeted Medicine, Breisacherstr. 113, 79106 Freiburg, Germany. equal contribution (Tilman Busch, Björn Neubauer).

## Abstract

Autosomal dominant polycystic kidney disease (ADPKD) is caused by mutations in *PKD1* and *PKD2*, encoding polycystin-1 (PC1) and polycystin-2 (PC2), which are required for the regulation of the renal tubular diameter. Loss of polycystin function results in cyst formation. Atypical forms of ADPKD are caused by mutations in genes encoding endoplasmic reticulum (ER)-resident proteins through mechanisms that are not well understood. Here, we investigate the function of DNAJB11, an ER co-chaperone associated with atypical ADPKD. We generated mouse models with constitutive and conditional *Dnajb11* inactivation and *Dnajb11*-deficient renal epithelial cells to investigate the mechanism underlying autosomal dominant inheritance, the specific cell types driving cyst formation, and molecular mechanisms underlying DNAJB11-dependent polycystic kidney disease. We show that biallelic loss of *Dnajb11* causes cystic kidney disease and fibrosis, mirroring human disease characteristics. In contrast to classical ADPKD, cysts predominantly originate from proximal tubules. Cyst formation begins *in utero* and the timing of *Dnajb11* inactivation strongly influences disease severity. Furthermore, we identify impaired PC1 cleavage as a potential mechanism underlying DNAJB11-dependent cyst formation. Proteomic analysis of *Dnajb11*- and *Pkd1*-deficient cells reveals common and distinct pathways and dysregulated proteins, providing a foundation to better understand phenotypic differences between different forms of ADPKD.

## Introduction

Polycystic Kidney Disease (PKD) represents a group of hereditary disorders characterized by the progressive development of fluid-filled cysts in the kidneys, leading to renal dysfunction and ultimately, end-stage kidney disease (ESKD) (1). Autosomal dominant inheritance is the prevailing genetic mechanism underlying PKD. Loss-of-function mutations in the genes *PKD1* and *PKD2* account for the majority of autosomal dominant polycystic kidney disease (ADPKD) (2, 3). The respective gene products polycystin-1 (PC1) and polycystin-2 (PC2) interact and form a receptor-ion channel complex that regulates renal tubular diameter and prevents cyst formation (4, 5). The folding and assembly of these large membrane proteins require the assistance of molecular chaperones and glycosylating proteins in the endoplasmic reticulum (ER) (6, 7). Atypical forms of ADPKD are caused by mutations in genes encoding ER-resident proteins including GANAB, ALG5, ALG8, and ALG9 (8–12). Loss of function of the respective ER proteins impairs PC1 maturation and trafficking, which has been proposed as a mechanism causing cyst formation in these rare forms of ADPKD (13).

To better understand the molecular components involved in PC1 biogenesis and function, we performed an unbiased PC1 interaction proteomics screen. One of the top hits of this screen was the co-chaperone DNAJB11, which co-immunoprecipitated with PC1 (Fig. S1 and Table S1). Shortly after we had found that DNAJB11 interacts with PC1, monoallelic mutations in *DNAJB11* were found to cause an atypical form of ADPKD which sparked our interest to study the role of DNAJB11 in cyst formation (14).

At the molecular level, DNAJB11 has been characterized as a member of the HSP40-chaperone family. It consists of three domains: an N-terminal J-Domain, that is common to all HSP40 chaperones and mediates their binding to HSP70-chaperones, a substrate-binding domain, and a C-terminal dimerization domain (15). DNAJB11 resides in the ER where it acts as one of the main co-factors of the HSP70-chaperone BiP thereby contributing to folding and assembly of secretory proteins and membrane proteins (16, 17). The role of DNAJB11 *in vivo* is poorly understood.

DNAJB11 is ubiquitously expressed (www.proteinatlas.org/ENSG00000090520-DNAJB11/tissue), which raises the question why patients with mutations in *DNAJB11* have only polycystic kidneys and few additional apparent phenotypes in other organs. Kidney cysts in patients with DNAJB11-related kidney disease are smaller than in classical ADPKD, kidney size is not increased and interstitial fibrosis is more pronounced. Furthermore, the course of the disease is slower with a median age of onset of ESKD in the 8th decade of life (18).

To investigate the role of DNAJB11 in PC1 biogenesis and cyst formation, we generated mouse models with constitutive and conditional inactivation of *Dnajb11* as well as *Dnajb11*-deficient renal epithelial cells. We aimed at addressing three fundamental questions: 1.) What is the genetic mechanism for autosomal dominant inheritance? 2.) Which cells drive cyst formation? 3.) Can we identify molecular mechanisms causing DNAJB11-dependent cyst formation?

In the classical form of ADPKD, a second-hit model provides a broadly accepted explanation for dominant inheritance as well as the focal nature of cystogenesis (19). According to this model, cyst formation is initiated by loss-of-heterozygosity due to inactivating somatic mutations in addition to the germline mutation in *PKD1* or *PKD2* (20). Cysts arise from homozygous mutant cells in all tubule segments if the function of PC1/PC2 decreases below a critical threshold well below 50% (13, 21). The precise mechanism how the PC1-PC2 complex regulates tubular morphology is still unknown. It is known, however, that PC1 undergoes autoproteolytic cleavage at its G protein-coupled receptor proteolytic site (GPS) with the two resulting fragments remaining non-covalently attached to each other (22). Mouse models expressing GPS-cleavage-deficient mutant PC1 develop kidney cysts highlighting the physiological importance of this posttranslational modification (23, 24).

In this study, we show that biallelic loss of *Dnajb11* in mice causes cystic kidney disease and fibrosis similar to the human disease. Mice with monoallelic inactivation of *Dnajb11* show no apparent phenotype establishing a cellular recessive mechanism of cyst formation. We find that cyst formation begins *in utero,* cysts originate almost exclusively from proximal tubules, and that postnatal conditional inactivation of *Dnajb11* strongly attenuates disease severity compared to constitutive inactivation. Finally, we show that loss of DNAJB11 strongly impairs GPS cleavage of PC1 *in vitro* and *in vivo* suggesting a mechanism of cyst formation involving PC1.

## Results

### Biallelic constitutive loss of *Dnajb11* causes cystic kidney disease

To investigate *Dnajb11* expression and function *in vivo*, we generated several *Dnajb11* mouse models based on a publicly available allele from the European Conditional Mouse Mutagenesis Program (EUCOMM) (25). We performed X-Gal stainings in *Dnajb11^tm1a/+^* mice harboring a LacZ cassette to monitor *Dnajb11* expression. At E10.5, *Dnajb11* expression was detected ubiquitously in the whole embryo (Fig. S2A). In the developing kidney *Dnajb11* expression was visible in all cell types at all investigated time points (Fig. S2B-D).

After removing the LacZ cassette, the resulting conditional floxed allele (*Dnajb11^fl^*) was converted into a constitutive knockout allele (*Dnajb11^−^*). To generate homozygous *Dnajb11^−/−^* mice, heterozygous *Dnajb11^+/−^* animals were intercrossed. The knockout was confirmed by absence of DNAJB11 protein in kidney lysates of *Dnajb11^−/−^* animals (Fig. 1F). Homozygous mice exhibited significant lethality with incomplete penetrance. Out of 61 weaned pups, only 7 (11%) were homozygous, much less than the expected Mendelian ratio (Fig. 1A). Two of the seven *Dnajb11^−/−^* mice died at five weeks of age of unknown cause. The five surviving *Dnajb11^−/−^ animals* displayed slightly, albeit not significantly reduced body weight at P90 (WT 23.2g ± 0.83 vs *Dnajb11^−/−^* 19.8g ± 2.26, p=0.20, n=5 each). In embryos at E17.5, the number of homozygous animals was slightly below the expected Mendelian ratio (17%) (Fig. 1C). All homozygous embryos had polycystic kidneys at E17.5 whereas kidneys of heterozygous embryos exhibited no pathologic phenotype (Fig. 1D). Homozygous embryos displayed no obvious extrarenal phenotypes.

**Fig. 1.**
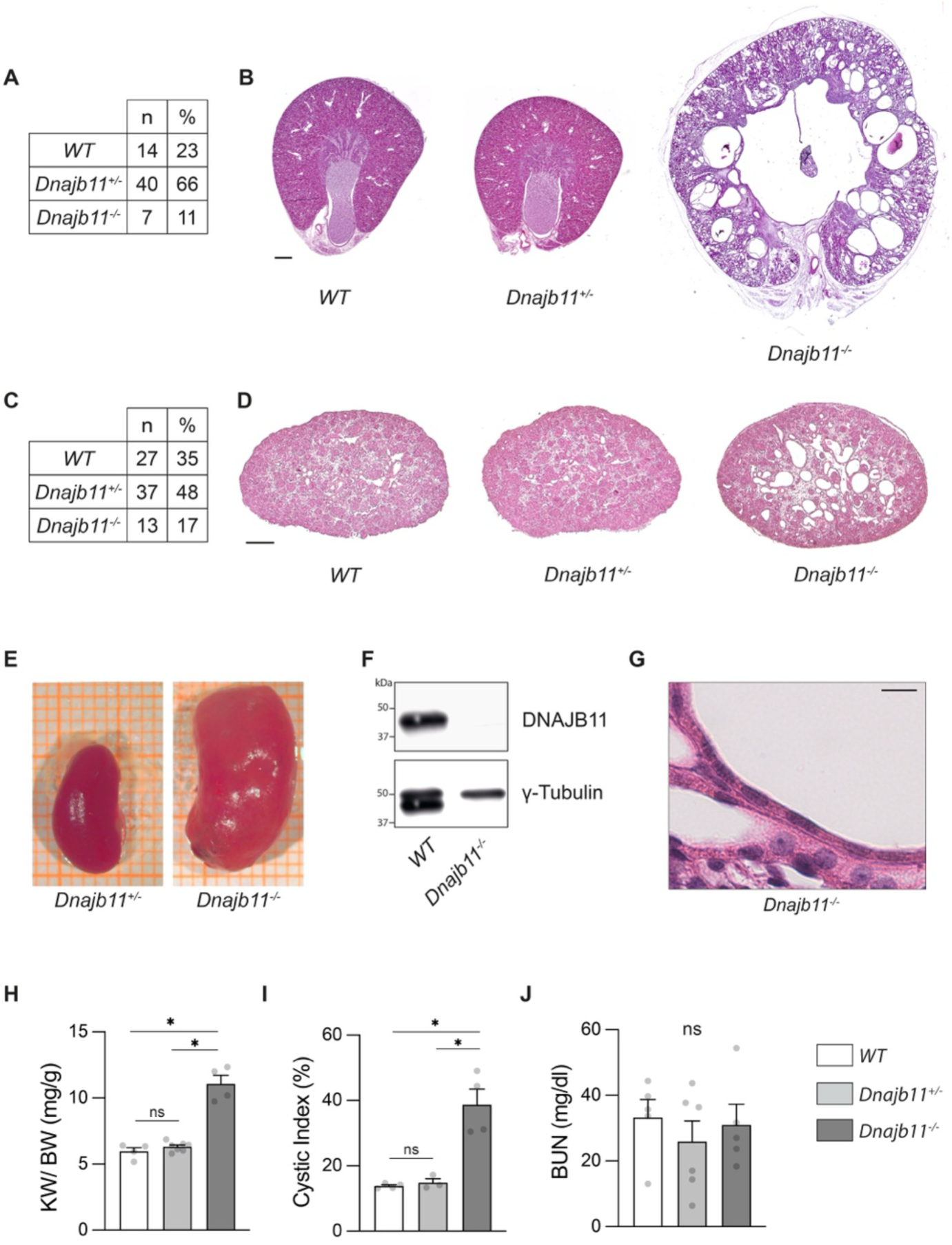
Constitutive inactivation of *Dnajb11*. In order to generate *Dnajb11^−/−^* mice, *Dnajb11^+/−^* animals were intercrossed. **(A), (C)** Distribution of genotypes of progeny from *Dnajb11^+/−^* intercrosses at P90 (A) and at E17.5 (C). Numbers of *Dnajb11^−/−^* progeny were lower than expected according to Mendelian ratios. **(B), (D)** HE stainings of kidney sections of P90 (B) and E17.5 (D) kidneys. No morphological abnormalities were observed in *Dnajb11^+/−^* kidneys. *Dnajb11^−/−^* kidneys displayed a cystic phenotype. Scale bar: 500 µm (B); 200 µm (D). **(E)** Macroscopic images of kidneys of *Dnajb11^+/−^* and *Dnajb11^−/−^* mice at P90. **(F)** DNAJB11 expression detected by anti-DNAJB11 immunoblotting of P90 kidney lysates. Dnajb11 is not detectable in lysates of *Dnajb11*^−/−^ kidneys. **(G)** High magnification of flattened cyst-lining epithelial cells. Scale bar: 20µm. **(H), (I), (J)** Aggregate quantitative data for KW/BW, cystic Index and BUN. KW/BW and cystic Index were significantly increased in *Dnajb11^−/−^* mice compared to WT and *Dnajb1^+/−^* mice. No statistically significant difference was observed in BUN between WT and *Dnajb11^+/−^* and WT and *Dnajb11^−/−^* mice. Bars represent mean of all replicates. Error bars indicate SEM. Statistical significance was evaluated using unpaired t-test. * indicates p<0.05. ns = non-significant.

Adult *Dnajb11^−/−^* animals (P90) developed enlarged polycystic kidneys with significantly increased cystic index and kidney weight/body weight (KW/BW) ratio, whereas *Dnajb11^+/−^* animals displayed no kidney pathology (Fig. 1B, E, H, I). Cyst formation was focal despite ubiquitous loss of DNAJB11 (Fig. 1B, D). The cyst lining epithelial cells showed a flattened cell shape as described for classical ADPKD (26, 27) (Fig. 1G). Despite these histological abnormalities, kidney function monitored by blood urea nitrogen (BUN) was not yet impaired in *Dnajb11^−/−^* animals compared to wild type (WT) at P90 (Fig. 1J). Adult homozygous *Dnajb11^−/−^* mice showed no obvious extrarenal phenotypes.

### Loss of DNAJB11 in the distal nephron does not cause kidney cysts

To circumvent the observed late embryonic or perinatal lethality and to further investigate the cellular origin of kidney cysts in *Dnajb11^−/−^* animals, we next used *Ksp-Cre* to selectively inactivate *Dnajb11* in tubular segments from the thick ascending loop of Henle to the collecting duct (28). *Ksp-Cre*-mediated deletion of *Pkd1* leads to the rapid development of a severe cystic phenotype (29). Surprisingly, *Ksp-Cre*-mediated deletion of *Dnajb11* did not lead to the development of a cystic kidney phenotype after one year (Fig. 2A, B). KW/BW, cystic index and BUN did not differ comparing *Ksp-Cre*; *Dnajb11^fl/−^* animals and the corresponding littermate controls (Fig. 2C-E).

**Fig. 2.**
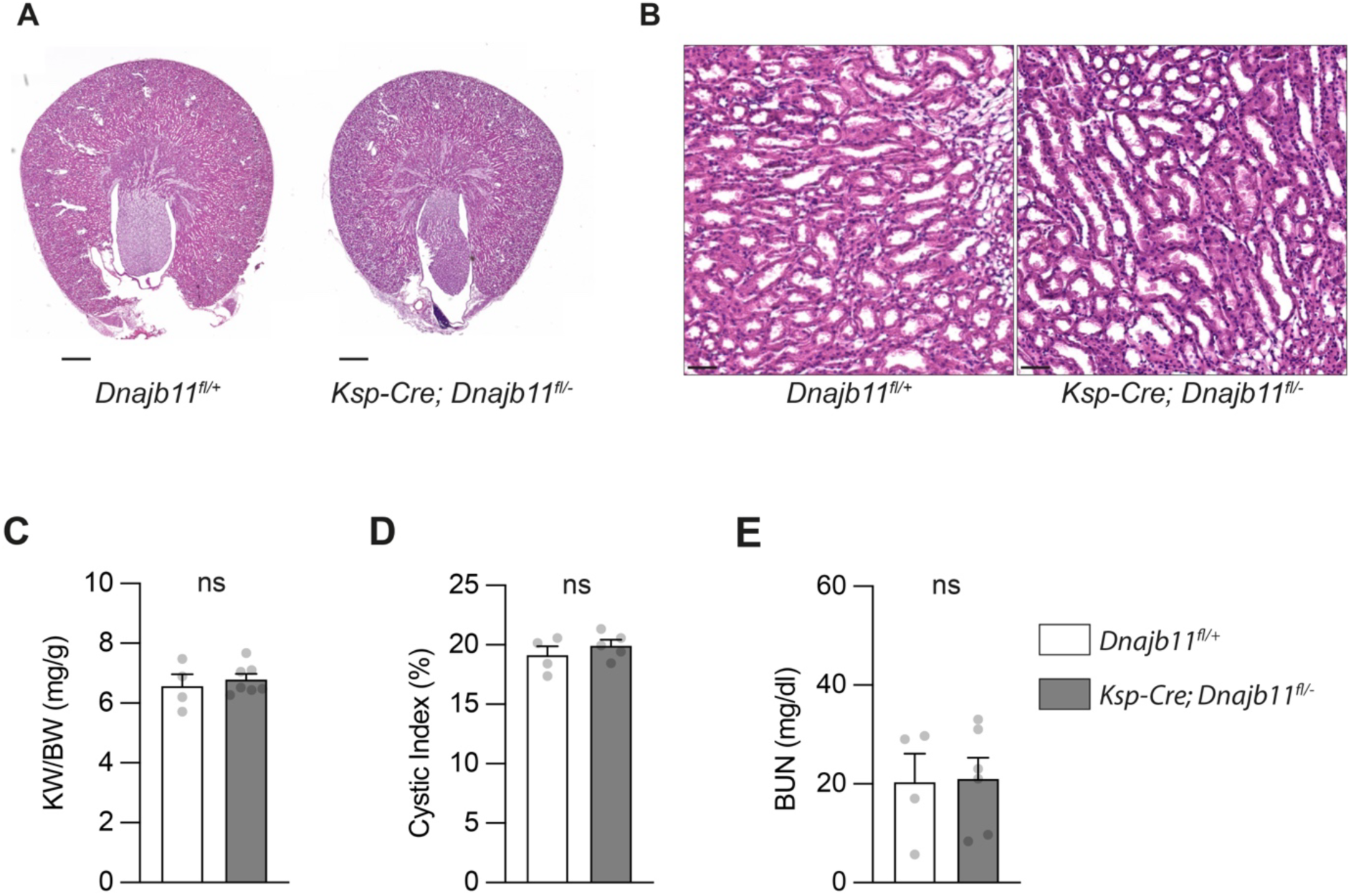
Distal tubule and collecting duct specific inactivation of *Dnajb11.* **(A), (B)** HE stainings of kidney sections at P360. *KspCre; Dnajb11^fl/−^* kidneys display no morphological abnormalities. Scale Bar: 500 µm (A), 50 µm (B). **(C), (D), (E)** Aggregate quantitative data for KW/BW, dilatation index, BUN. No significant differences were observed between experimental groups. Bars represent mean of all replicates. Error bars indicate SEM. Statistical significance was evaluated using unpaired t-test. ns = non-significant (p>0.05).

### Kidney cysts in *Dnajb11^−/−^* mice originate from the proximal tubule

To investigate the reason for the absence of a kidney phenotype in *Ksp-Cre*; *Dnajb11^fl/−^* animals, we assessed which nephron segments are the origin of cysts in constitutive *Dnajb11^−/−^* mice. Using markers of different nephron segments, we found that cyst-lining epithelial cells in *Dnajb11^−/−^* kidneys at E17.5 were virtually all positive for Low Density Lipoprotein receptor-related Protein 2 (LRP2), a marker of the proximal tubule (Fig. 3A) (30); (31). None of the cysts in embryonic kidneys were positive for markers of other nephron segments (Fig. 3A, Fig. S3A, B). Similarly, at P90 the vast majority of cysts were also positive for LRP2 with a few slightly cystic tubules with calbindin staining (CALB) (Fig. 3B), a marker of the distal tubule (32). Like in embryonic kidneys, cyst epithelial cells were not labeled by antibodies against uromodulin (UMOD) and aquaporin-2 (AQP2), which are marker proteins for the thick ascending loop of Henle and the collecting duct, respectively (Fig. S3C) (33, 34). Taken together, cysts in *Dnajb11^−/−^* kidneys originate from the proximal tubule, thus explaining the absence of kidney cysts in *Ksp-Cre*; *Dnajb11^fl/−^* animals.

**Fig. 3.**
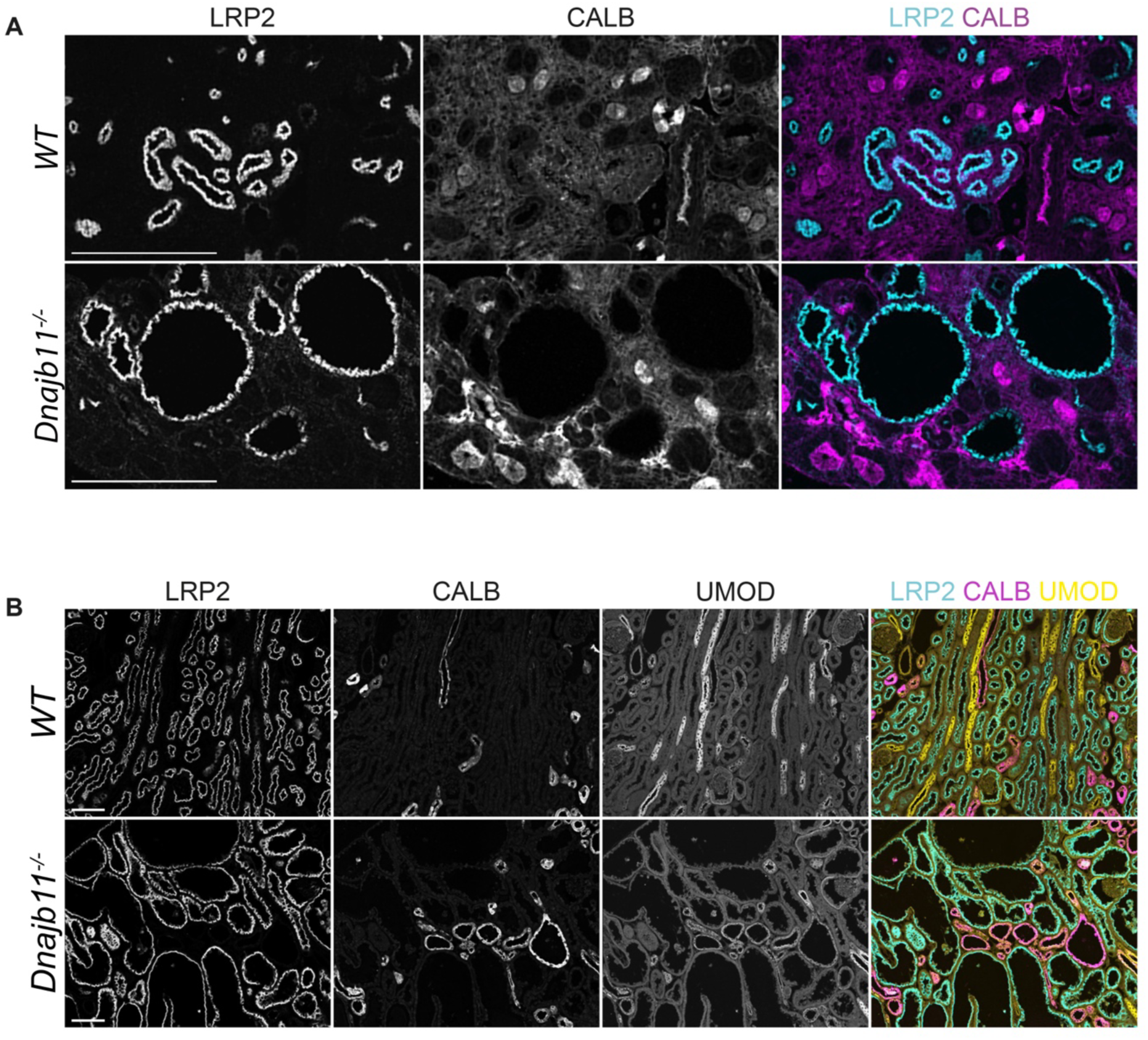
Immunohistochemistry using tubule segment markers. **(A), (B)** E17.5 (A) and P90 (B) WT and *Dnajb11^−/−^ k*idney sections were labeled with kidney tubule segment markers LRP2 and CALB. At E17.5, cyst-lining epithelium is exclusively labeled by LRP2. At P90, larger cysts are also exclusively labeled by LRP2. Few dilated tubules are positive for CALB. Scale Bar: 200 µm (A), 100 µm (B)

### Pan-tubular *Dnajb11* inactivation starting at P28 causes proximal tubular dilation

To confirm the proximal tubular origin of cysts in *Dnajb11^−/−^* kidneys, we opted for an alternative doxycycline-inducible Cre allele: *Pax8rtTA;LC1-Cre* (referred to as *Pax8-Cre*). Upon induction Cre-recombinase is active in the complete renal tubular system of these animals, including the proximal tubule. We induced *Pax8-Cre; Dnajb11^fl/fl^* mice from P28 to P42 since this induction protocol has been shown to cause a moderate to severe cystic phenotype 12-14 weeks after inactivation of *Pkd1* or *Pkd2* (35, 36). 12 weeks after induction *Pax8-Cre; Dnajb11^fl/fl^* mice had slightly larger kidneys with a significantly increased KW/BW ratio (Fig. 4A, C). The histology showed widespread tubular dilations resulting in a significantly increased dilation index compared to non-induced controls (Fig. 4B, D). These tubular dilations are known precursor stages of kidney cyst (27). Consistent with the mild phenotype, no significant difference in kidney function (BUN) was detected in induced mice compared to the control group (Fig. 4E). These experiments show that postnatal inactivation of *Dnajb11* strongly attenuates disease severity compared to constitutive inactivation. Dilated tubules in tubule-specific *Dnajb11^−/−^* knockout mice were exclusively positive for LRP2 (Fig. 4F). No dilated tubules were found to be positive for CALB, UMOD or for AQP2 (Fig. S4A, B). This supports the observation in constitutive *Dnajb11^−/−^* knockout animals that kidney cysts arise predominantly if not exclusively from proximal tubules.

**Fig. 4.**
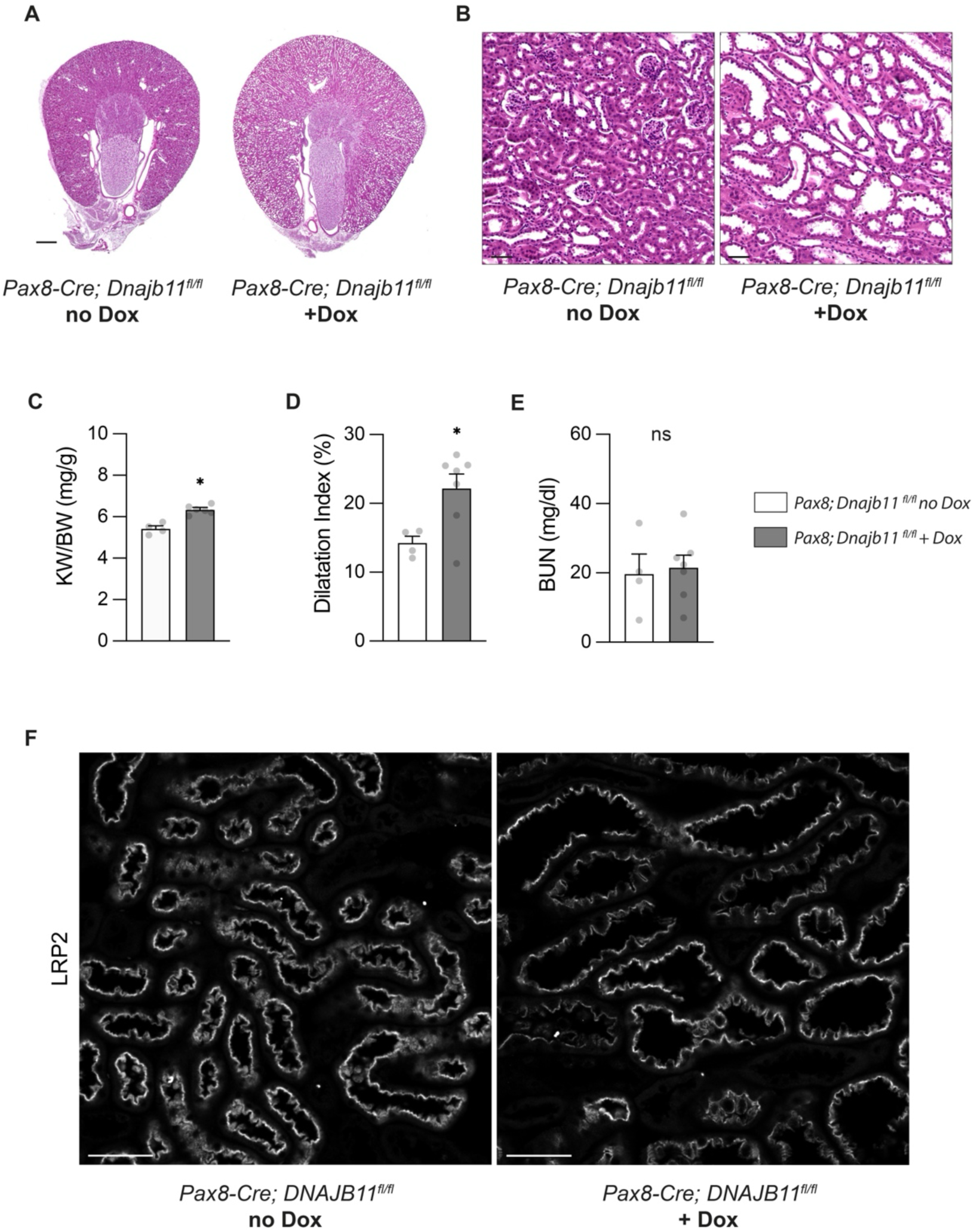
Pan-tubular inactivation of *Dnajb11.* **(A), (B)** HE stainings of kidney sections of uninduced (left) and doxycycline-induced (right) *Pax8-Cre; Dnajb11^fl/fl^* mice at P126; Kidneys from doxycycline-induced animals display tubular dilatations. Scale bar: 500 µm (A), 50 µm (B) **(C), (D), (E)** Aggregate quantitative data for KW/BW, dilation index and BUN; KW/BW and dilation index are significantly increased in Doxycyclin-induced *Pax8-Cre; Dnajb11^fl/fl^* mice versus non-induced controls. **(F)** Kidney sections of non-induced and doxycycline-induced Pax8-Cre; *Dnajb11^fl/fl^* mice were immunolabeled for proximal tubule marker LRP2. Dilated tubules are positive for LRP2. Bars represent mean of all replicates. Error bars indicate SEM. Statistical significance was evaluated using unpaired t-test. * indicates p<0.05. ns = non-significant.

### Fibrosis in *Dnajb11^−/−^* animals

Human *DNAJB11*-related kidney disease encompasses the development of extensive interstitial fibrosis in addition to kidney cysts. To investigate the development of fibrosis in our *Dnajb11*-deficient mouse models, we stained kidney sections with an antibody against α-smooth muscle actin (α-SMA). In kidney sections of constitutive *Dnajb11^−/−^* knockout animals, we found large, mainly pericystic fibrotic areas (Fig. 5A). Acid Fuchsin staining confirmed this finding (Fig. S5). In contrast, kidney sections of *Ksp-Cre; Dnajb11^fl/−^* mice did not show SMA-positive fibrotic areas (Fig. 5B) suggesting that fibrosis upon loss of *Dnajb11* does not occur independently of cystogenesis. Accordingly, *Pax8-Cre; Dnajb11^fl/fl^* mice, which display tubular dilations after postnatal induction but no prominent cysts, did not show signs of fibrosis (Fig. 5C). The fact that there are no discernible fibrotic changes prior to the development of kidney cysts in *Dnajb11*-deficient animals suggests that, at least in mice, fibrosis occurs secondary to cyst formation.

**Fig. 5.**
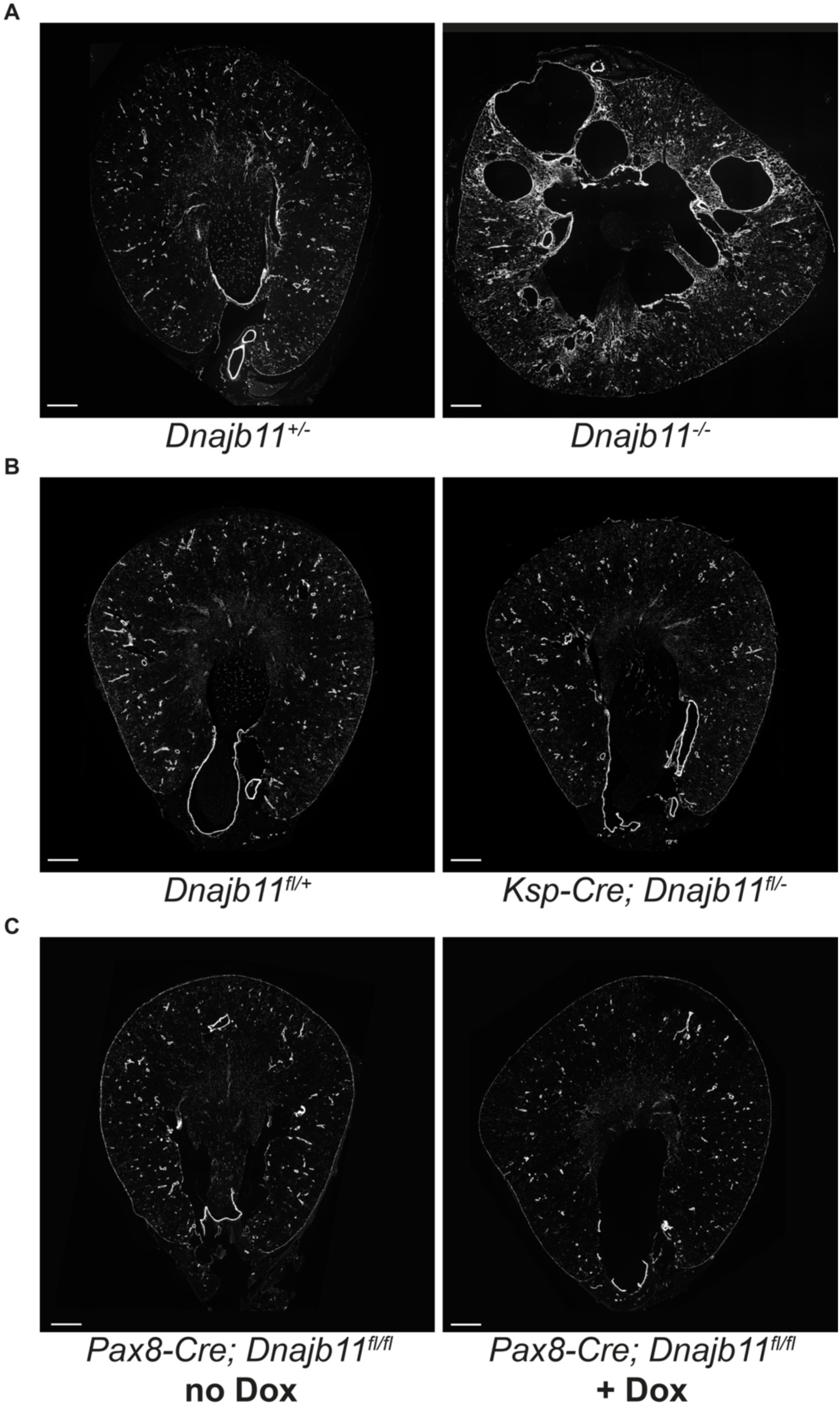
Increased fibrosis in *Dnajb11^−/−^* kidneys. **(A), (B), (C)** Immunofluorescence staining of SMA as a marker for fibrosis. **(A)** Fibrosis is increased in kidneys of P90 *Dnajb11^−/−^* mice compared to kidneys of *Dnajb11^+/−^*. **(B)** Fibrosis is not increased in kidneys of *KSP-Cre; Dnajb11^fl/−^* animals compared to controls at P360 as well as **(C)** in kidneys of doxycycline-induced *Pax8-Cre; Dnajb11^fl/−^* animals compared to non-induced controls at P126. Scale Bar: 500 µm

### PC1 cleavage is impaired in *Dnajb11^−/−^* cells and kidneys

Impaired PC1 processing has been implicated as a potential pathogenic mechanism in *Dnajb11*-related kidney disease(37). In order to investigate the molecular mechanism underlying *Dnajb11*-related kidney disease and whether the interaction between DNAJB11 and PC1 has an effect on the biogenesis and processing of PC1, we used CRISPR/Cas9 to generate *Dnajb11*-deficient cell lines. We deleted *Dnajb11* in WT IMCD3 cells, as well as in an IMCD3 cell line in which we had previously added N-terminal FLAG-tags and C-terminal V5-tags to endogenous PC1 via CRISPR/Cas9 and homologous recombination (Fig. 6A, B). In this cell line, endogenous full length (FL) PC1 as well as the cleaved C-terminal fragment (CTF) can be detected via the V5 epitope tag. After immunoprecipitation and immunoblotting with a V5 antibody, we observed a significant reduction in the ratio of cleaved PC1 CTF versus uncleaved FL PC1 (Fig. 6C, D). Notably, re-expression of human DNAJB11 shifted the CTF/FL ratio back towards normal (Fig. 6G, H).

**Fig. 6.**
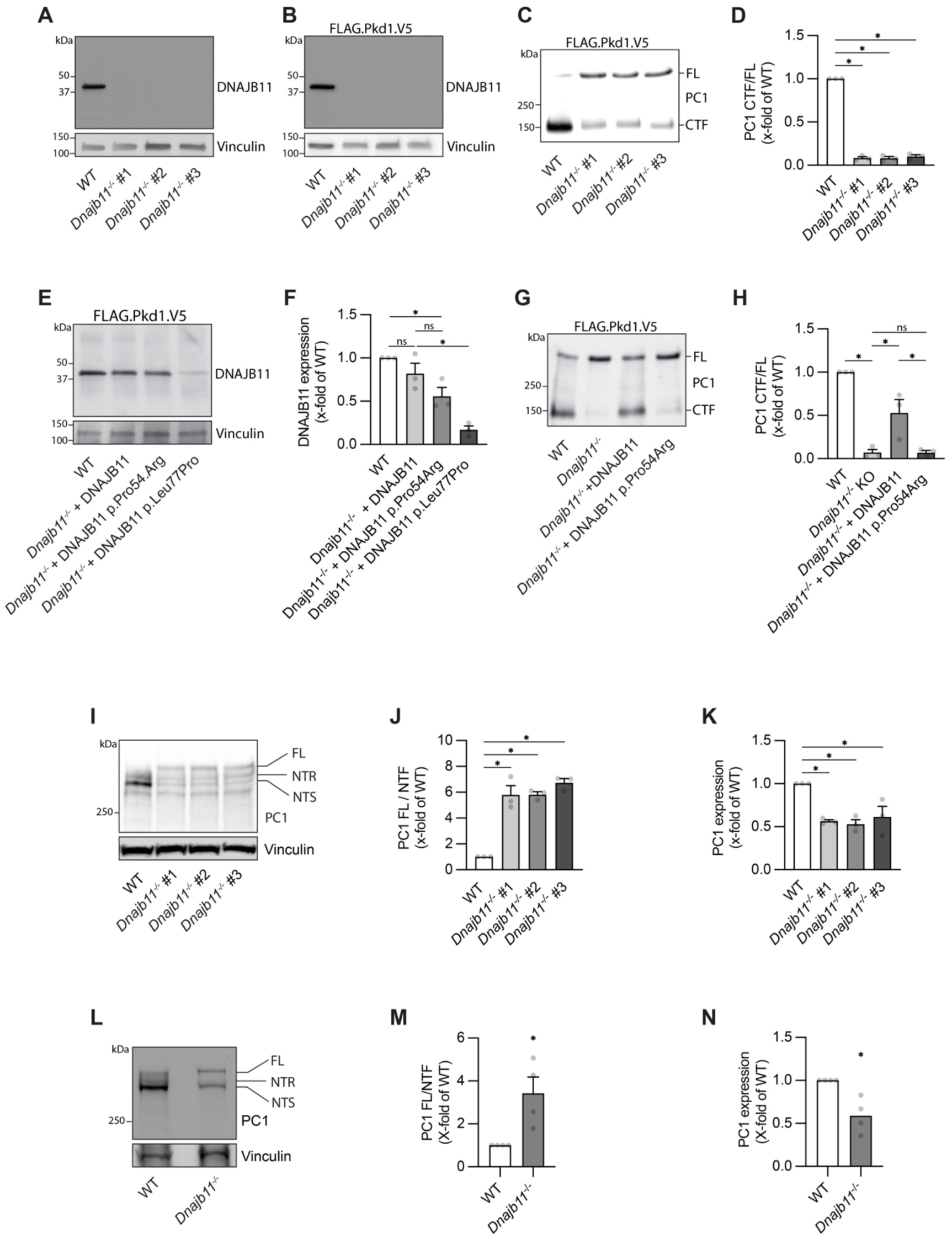
Loss of DNAJB11 leads to impaired PC1 processing *in vitro* and *in vivo*. **(A)** DNAJB11 expression detected by anti-DNAJB11 immunoblotting *in* IMCD WT and 3 IMCD *Dnajb11^−/−^ c*ell lines. Three independent clones of genome-edited cell lines were studied to exclude clonal artifacts. **(B)** DNAJB11 expression detected by anti-DNAJB11 immunoblotting in IMCD WT FLAG.Pkd1.V5 and IMCD FLAG.Pkd1.V5 *Dnajb11^−/−^* cells. **(C)** C-terminally V5-tagged PC1 was immunoprecipitated via Protein A coupled V5 antibody from lysates of IMCD WT Flag.Pkd1.V5 and 3 IMCD FLAG.Pkd1.V5 *Dnajb11^−/−^* cell lines. **(D)** Densitometric quantification of ratio of PC1 CTF/ PC1 FL. CTF/FL ratio is significantly decreased in all three IMCD FLAG.Pkd1.V5 *Dnajb11^−/−^* cell lines. **(E)** Reexpression of DNAJB11, DNAJB11 p.Pro54Arg and DNAJB11 p.Leu77Pro in IMCD FLAG.Pkd1.V5 *Dnajb11^−/−^*. **(F)** Densitometric quantification of DNAJB11 expression. **(G)** C-terminally V5-tagged PC1 was immunoprecipitated from lysates of IMCD WT, IMCD *Dnajb11^−/−^*, IMCD *Dnajb11^−/−^* + DNAJB11 WT and IMCD *Dnajb11^−/−^* + DNAJB11 p.Pro54Arg cell lines. **(H)** Densitometric quantification of ratio of PC1 CTF/ PC1 FL. CTF/FL ratio is rescued only by WT DNAJB11 but not by DNAJB11 p.Pro54Arg. **I** PC-1 expression detected by anti-PC1 (7E12) immunoblotting in IMCD WT and 3 IMCD *Dnajb11^−/−^* cell lines. **(J)**, **(K)** Densitometric quantification of ratio of (J) PC1 FL/NTF and (K) overall PC1 expression (FL + NTR + NTS) in IMCD WT and 3 IMCD *Dnajb11^−/−^* cell lines. **(L)** PC1 expression detected by anti-PC1 (7E12) immunoblotting in kidney lysates of WT and *Dnajb11^−/−^* mice (E17.5). **(M)**, **(N)** Densitometric quantification of ratio of (M) PC1 FL/ NTF. NTF = NTS+ NTR. (N) Densitometric quantification of overall PC1 expression (FL + NTR + NTS) in kidney lysates of WT and *Dnajb11^−/−^* mice (E17.5). Bars represent mean of all replicates. Error bars indicate SEM. Statistical significance was evaluated using unpaired t-test. * indicates p<0.05. ns = non-significant.

To investigate if disease-causing *Dnajb11* missense variants impair PC1 cleavage, we expressed two disease-causing DNAJB11 mutants in *Dnajb11^−/−^* cells (37). Mutations leading to singular amino acid substitutions rather than truncating mutations were chosen to test whether PC1 cleavage can serve as a biochemical readout for DNAJB11 variants of unknown significance. DNAJB11 p.Pro54Arg was expressed at levels similar to WT DNAJB11 in *Dnajb11^−/−^* cells whereas DNAJB11 p.Leu77Pro was barely detectable (Fig. 6E, F). We tested the well expressed DNAJB11 p.Pro54Arg mutant for its ability to rescue PC1 cleavage. Quantification of the PC1 CTF/FL ratio shows that expression of DNAJB11 p.Pro54Arg cannot rescue the PC1 cleavage defect in *Dnajb11*^−/−^ cells despite its robust expression, thus demonstrating that disease-causing DNAJB11 mutations impair PC1 cleavage (Fig. 6 G, H).

The impairment of PC1 cleavage could be confirmed in non-tagged *Dnajb11^−/−^* cells. Immunoblotting and detection of PC1 with an N-terminal PC1 antibody shows an increase in the relative amount of uncleaved FL PC1 in comparison to the two cleaved N-terminal fragments (NTF); EndoH-resistant (NTR) and EndoH-sensitive (NTS) N-terminal PC1. Furthermore, the overall amount of PC1 in *Dnajb11^−/−^* cells was decreased (Fig. 6I, K). Finally, we confirmed the impairment of PC1 cleavage in kidney lysates of mouse embryos at E17.5. In *Dnajb11^−/−^* mice, there is a significant increase in the ratio of PC1 FL/NTF compared to WT controls (Fig. 6L, M). In addition, the decrease in the overall amount of PC1 observed *in vitro* could also be confirmed in kidney lysates of *Dnajb11^−/−^* mice (Fig. 6L, N).

### Proteomics of *Dnajb11*- and *Pkd1*-deficient renal epithelial cells

DNAJB11 has been described as one of the main co-factors of the HSP70-chaperone BiP and contributes to the biogenesis and processing of proteins. The impaired maturation and cleavage of PC1 in *Dnajb11*-deficient cells supports this notion and represents a plausible mechanism contributing to cyst formation. Yet, the different clinical presentation of ADPKD caused by loss of either DNAJB11 or PC1 raises the question whether there are also distinct pathogenic mechanisms for each disease. To gain insights into additional client proteins of DNAJB11, we performed quantitative proteomics in WT versus DNAJB11-deficient IMCD3 cells (Table S2). As obvious candidates, we looked for downregulated proteins encoded by genes causing cystic disease and found no significant differences (Fig. 7F).

**Fig. 7.**
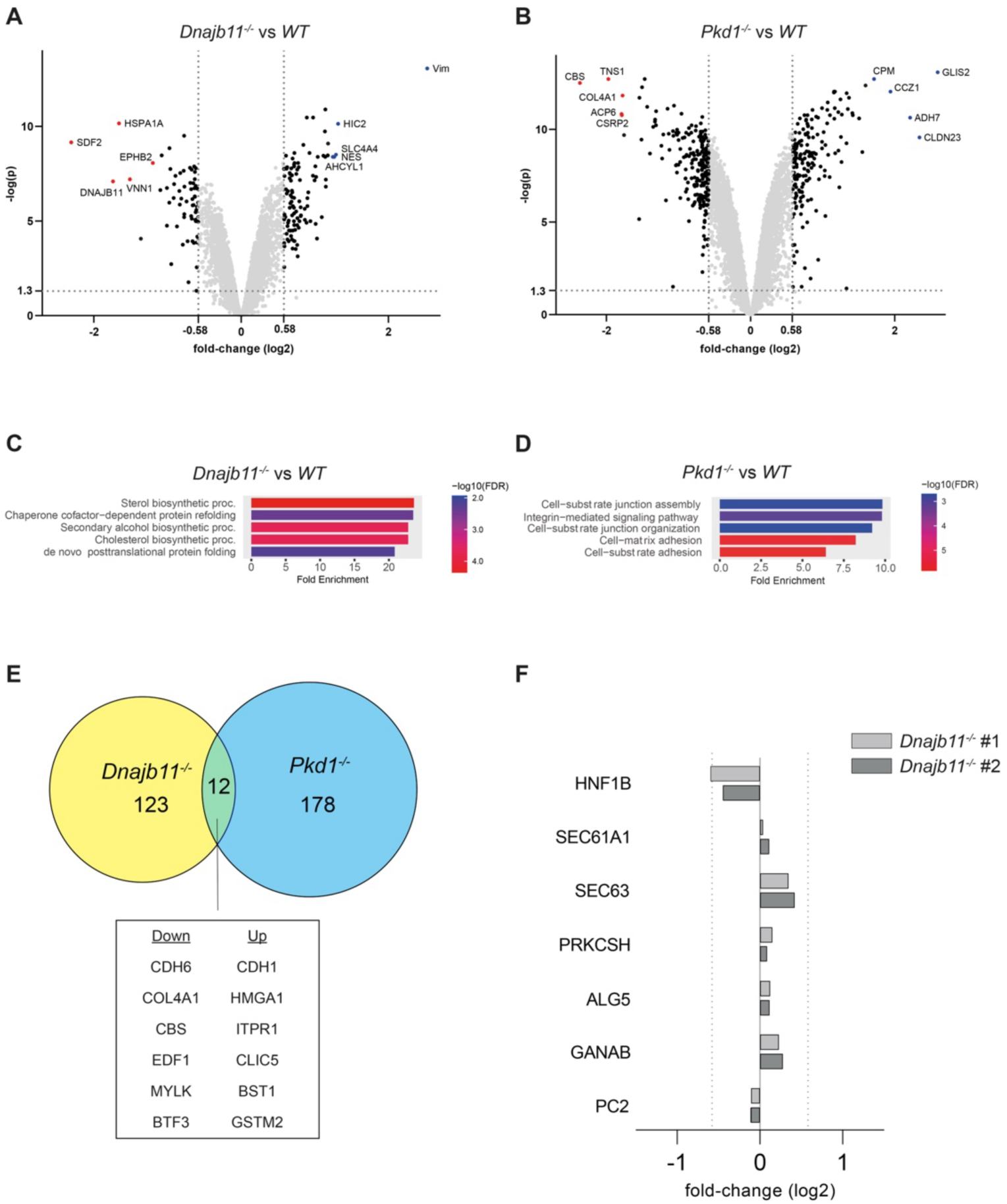
Proteome analysis of DNAJB11- and PC1-defecient renal epithelial cells. **(A), (B)** Volcano plot indicating differentially expressed proteins between (A) IMCD *DNAJB11^−/−^* vs WT and (B) IMCD3 *Pkd1^−/−^* vs WT cells. Annotated are the 5 proteins with the highest fold-enrichment that were detected in both investigated *Dnajb11^−/−^* clones (A) and in both investigated Pkd1^−/−^ clones (B). Threshold for adjusted p-value was set to log_10_ > 1.3. Threshold for fold-change was set to log_2_ +/− 0.58. **(C), (D)** GO term analysis of differentially regulated proteins in *Dnajb11^−/−^* vs WT and *Pkd1^−/−^* vs WT cells. **(E)** Venn-diagram illustrating the overlap of differentially regulated proteins between *Dnajb11^−/−^* and *Pkd1^−/−^* cells. 123 proteins were significantly and concordantly differentially expressed in both *Dnajb11^−/−^* clones, 178 proteins were significantly and concordantly differentially expressed in both *Pkd1^−/−^* clones. 12 proteins are concordantly up-/downregulated in *Dnajb11^−/−^* and *Pkd1^−/−^* cells compared to WT cells. **(F)** Expression levels of known cyst genes in *Dnajb11^−/−^* clones #1 and #2 compared to WT cells. Threshold for fold-change was set to log_2_ > 0.58 or log_2_ < −0.58.

In a more unbiased discovery-based approach we compared similarities and differences between DNAJB11- and PC1-deficient cells. These experiments showed that loss of both proteins changed the expression levels of many proteins significantly (Fig. 7A, B; Tables S2-4). Pathway enrichment analyses showed enrichment of distinct pathways for loss of DNAJB11 and PC1 (Fig. 7C, D). While the Top 5-enriched pathways highlighted a role for DNAJB11 in sterol/cholesterol biosynthesis and chaperone-dependent protein folding, PC1-dependent pathways included cell substrate and junction assembly, integrin-mediated signaling and cell-matrix adhesion. At the level of individual proteins, the overlap of differentially regulated proteins was rather limited (Fig. 7E, Table S2-4). Out of 123 differentially expressed proteins in DNAJB11-deficient and 178 in PC1-deficient cells, only 12 showed significant concordant differential regulation in both knockout cell lines, despite the fact that both knockouts were generated from the same isogenic parental cell line (38). These commonly regulated proteins may provide insights into shared pathogenic mechanisms of cyst formation and deserve further investigation. For example, mutations to COL4A1, which is downregulated in both cell lines, have been proposed to cause an atypical form of ADPKD, raising the possibility of a common mechanistic role in cyst formation in different forms of ADPKD (39, 40).

In addition to commonalities, differences between DNAJB11-deficient and PC1-deficient cells are also of interest. To better understand the distinct clinical features of DNAJB11-related kidney disease differentially regulated proteins that are only found in DNAJB11-deficient cells may provide interesting entry points for further studies. Such proteins may help to identify other disease-relevant client proteins as well as additional co-factors in the chaperone network contributing to the structural integrity of kidney tubules. Of note, the most strongly downregulated protein in DNAJB11-deficient cells, SDF2, has recently been described as an interactor of DNAJB11 regulating its chaperone function(41). Further studies are required to unravel the precise role of such candidate proteins in DNAJB11 function.

## Discussion

In this study, we show that homozygous deletion of *Dnajb11* causes a kidney phenotype that recapitulates essential aspects of human *DNAJB11*-related kidney disease and provides insights into pathogenic mechanisms. While monoallelic mutations in *DNAJB11* are sufficient to cause kidney disease in humans, heterozygous *Dnajb11^+/−^* mice do not develop kidney cysts. Inactivation of both *Dnajb11* alleles causes cyst formation showing that *Dnajb11*-related kidney disease is recessive at the cellular level in mice. If this genetic mechanism applies to humans, cyst formation would require somatic *DNAJB11* mutations resulting in loss-of-heterozygosity. Future studies in human samples from patients with cystic disease caused by monoallelic *DNAJB11* mutations will have to test whether cyst epithelia indeed harbor somatic mutations as reported for *PKD1* or *PKD2*(20). As shown recently, cyst burden and kidney enlargement in *Dnajb11*-deficient mice are much milder than in *Pkd1-* or *Pkd2-*deficient mouse models (42). This corresponds with the human disease, which is also characterized by a milder cystic phenotype and slower disease progression compared to classical ADPKD (18).

Knowledge about the cellular origin of cyst formation is critical to understand the underlying pathogenic mechanisms. Surprisingly, we find that cysts in *Dnajb11*^−/−^ mice originate from proximal tubules. The unexpected finding that *Ksp-Cre*-mediated inactivation of *Dnajb11* does not result in cystic kidneys suggests that cysts are not derived from distal tubular segments where *Ksp-Cre* is expressed (thick ascending loop of Henle, distal tubule, and collecting duct). A recent study also reported the absence of cysts in *Ksp-Cre; Dnajb11^fl/fl^* mice (42). It was shown in this study that a reduction in *Pkd1* copy number (*Pkd1^+/−^*) in these animals resulted in cyst formation (42). Since patients with ADPKD caused by monoallelic *DNAJB11* mutations do not have heterozygous *PKD1* mutations it was speculated that a lower basal PC1 dosage in humans compared to mice may explain cyst formation in the human disease (42). Our finding that cysts in *Dnajb11*-deficient mice are derived from proximal tubules but not from distal tubule segments provides a simple explanation for cyst formation without the requirement of additional factors or genetic manipulations. The proximal tubular origin of cysts is supported by three observations: 1.) Cysts in constitutive *Dnajb11^−/−^* kidneys are positive for LRP2; 2.) *Pax-Cre; Dnajb11^−/−^* mice show cystic dilations that are exclusively marked by LRP2; and 3.) *Ksp-Cre*-mediated *Dnajb11* inactivation does not cause cyst formation. Since expression of the *Ksp-Cre* transgene starts in mid-embryogenesis (28), one could speculate that earlier inactivation of *Dnajb11* in distal tubule segments might cause cyst formation. Yet, the extensive proximal tubular dilation in *Pax-Cre; Dnajb11^−/−^* mice after postnatal induction (P28-P42) argues against this. We occasionally observed small cysts that were positive for CALB in adult constitutive *Dnajb11^−/−^* animals in advanced stages of cystic disease. However, we never observed CALB-positive cysts in embryos or in *Pax-Cre; Dnajb11^−/−^* mice. The rare occurrence of CALB-positive small cysts at later time points may either be secondary due to strongly altered tissue architecture in advanced stages of cystic transformation or due to cell-autonomous effects giving rise to cyst formation in a few CALB-positive cells. Taken together, our data suggest that kidney cysts resulting from loss of DNAJB11 arise predominantly if not exclusively from proximal tubules. This is different in ADPKD caused by *PKD1* or *PKD2* mutations where cyst formation occurs in all nephron segments (43, 44). The intriguing question why cysts form only in proximal tubules remains to be clarified. Possible explanations include differential requirements for PC1 expression and GPS cleavage in different nephron segments or yet to be identified proteins with segment-specific roles in the regulation of tubular morphology. The knowledge about the cellular origin of cysts in *DNAJB11*-related ADPKD is important for future studies investigating mechanisms of cyst formation *in vivo* and may have implications for therapeutic interventions with drugs such as Tolvaptan which acts on Vasopressin 2 receptors that are not expressed in the proximal tubule.

The timing of inactivation of *Pkd1* or *Pkd2* strongly affects the severity of the cystic phenotype in mice. Conditional inactivation of *Pkd1* after P14 results in a much milder phenotype than inactivation before P13 (45). To test whether the timing of inactivation also affects the phenotype in *Dnajb11*-deficient animals, we used doxycycline-inducible *Pax-Cre; Dnajb11^fl/fl^* mice to delete *Dnajb11* in all nephron segments. Induction from P28 to P42 resulted in widespread proximal tubular dilations at P126. This phenotype was much milder than the phenotype of constitutive *Dnajb11*-deficient animals at P90 demonstrating that disease severity is indeed affected by the timing of *Dnajb11* inactivation.

The phenotypic spectrum of patients with mutations in *DNAJB11* is milder than in patients with *PKD1* mutations. Our mouse models recapitulate this observation. Both constitutive and inducible *Dnajb11* deletion with *Pax8-Cre* result in less severe phenotypes than the respective *Pkd1*-deficient mouse models (36), corroborating phenotypic differences caused by loss of the respective ADPKD genes. Furthermore, the likelihood of somatic mutations causing loss of heterozygosity is much smaller in patients with *DNAJB11* germline mutations than in patients with *PKD1* mutations, a gene with a much larger coding sequence. Notably, bi-allelic mutations in *DNAJB11* have recently been reported to cause Ivemark II syndrome, a severe fetal disorder with polycystic kidneys as well as pancreas and liver abnormalities (46). Homozygous loss of *DNAJB11* causes enlarged cystic kidneys in embryos like in our constitutive knockout mouse. Surprisingly, a recent study by Roy et al. did not find cyst formation in embryonic kidneys of constitutive *Dnajb11^−/−^* mice (42). The fact that cystic kidneys in patients with monoallelic *DNAJB11* mutations are not enlarged may be explained by the fact that focal somatic mutations in a small subset of tubular epithelial cells cause a lower cyst burden than complete loss of function in all cells. Our observation that the timing of inactivation strongly affects disease severity in mice may also contribute to less severe phenotypes since somatic mutations occurring later in life probably result in less prominent cyst formation and progression in patients.

Considering the proposed fundamental role of DNAJB11 in protein biogenesis and its ubiquitous expression, the observed phenotypes can be considered surprisingly mild (16, 17). We find that constitutive homozygous deletion of *Dnajb11* in mice leads to reduced viability. The number of homozygous *Dnajb11^−/−^* knockout animals was already lower than expected according to Mendelian ratios in late embryonic development (E17.5) and even further reduced after birth. The rather mild kidney phenotype cannot explain the lethality. Yet, homozygous *Dnajb11^−/−^* mice displayed no obvious extrarenal phenotypes. Further studies will be needed to address the reason for the reduced viability of *Dnajb11^−/−^* mice.

Fibrosis is a prominent feature in patients with monoallelic *DNAJB11* mutations. The kidney phenotype has been reported to be between the spectrum of ADPKD and autosomal dominant tubulo-interstitial kidney disease (ADTKD). It was therefore proposed that the mechanism causing interstitial fibrosis may involve ADTKD gene products such as uromodulin in the thick ascending loop of Henle (TAL). This hypothesis was supported by the observation of a possible intracellular accumulation of uromodulin in two kidney biopsies from patients with heterozygous *DNAJB11* mutations (14). We found prominent fibrosis in kidneys of *Dnajb11^−/−^* mice, mainly in areas surrounding cysts. However, we did not find obvious differences between uromodulin staining patterns in WT and *Dnajb11*-deficient animals (Fig. 3A, Fig. S4). This finding is supported by a recent study questioning the involvement of uromodulin in *DNAJB11*-dependent kidney disease based on data measuring urinary uromodulin levels and its trafficking in renal epithelial cells (47). Furthermore, we did not find evidence for an involvement of the TAL in cyst formation or fibrosis. *Ksp-Cre*-mediated *Dnajb11* inactivation in distal tubule segments including the TAL did not lead to cyst formation or fibrosis, even in 12 months old animals. Interestingly, kidney cysts in Ivemark II syndrome were also found to be exclusively uromodulin-negative (46). Our finding that pan-tubular inactivation of *Dnajb11* leads to proximal tubular dilatations without fibrosis, together with the above-mentioned observations, suggests that fibrosis in *Dnajb11*-deficient animals occurs as a consequence of cyst formation. It is possible that the extensive interstitial fibrosis in patients with *DNAJB11* mutations develops over long periods of time that cannot be investigated in the much shorter life span of mice. The mechanisms leading to extensive fibrosis remain to be determined and may involve inflammatory processes and yet to be determined client proteins of DNAJB11 (35, 48).

Impaired PC1 processing and maturation has been implicated as a potential pathogenic mechanism in *DNAJB11*-related kidney disease (37). Indeed, we find that GPS-cleavage of PC1 is strongly impaired after homozygous loss of *Dnajb11*. This robust biochemical phenotype is found in cell culture and *in vivo,* confirming recent findings (42). Furthermore, we demonstrate that DNAJB11 with a disease-causing mutation resulting in a single amino acid substitution leads to impaired PC1 processing, even though the mutated protein is expressed at levels similar to WT DNAJB11 (Fig. 6 E-H). This suggests, that PC1 GPS cleavage can be used to test the pathogenicity of *DNAJB11* missense mutations of unknown significance. The physiological relevance of PC1 cleavage is well established. Two mouse models expressing GPS-cleavage-deficient PC1 have been shown to develop polycystic kidneys, with one model behaving like a complete null allele and another model showing hypomorphic effects with a milder phenotype than a germline *Pkd1* knockout (23, 24). Yet, the phenotype of both cleavage-deficient mouse models was more severe than the phenotype of *Dnajb11^−/−^* animals. Possible explanations for these phenotypic differences include: 1.) PC1 cleavage is strongly reduced but not completely eliminated after loss of *Dnajb11,* both *in vitro* and *in vivo*. Since a PC1 dosage of about 20% has been shown to be sufficient to prevent cystogenesis (49), the small remaining fraction of cleaved PC1 may be sufficient to prevent more severe cystogenesis in kidneys of *Dnajb11^−/−^* mice. 2.) Cystogenesis caused by loss of *Dnajb11* is limited to proximal tubules which may contribute to a milder phenotype than reduced PC1 function which also involves distal tubule segments.

Impaired PC1 cleavage likely contributes to cyst formation upon loss of DNAJB11. Nevertheless, it seems prudent to investigate additional pathogenic mechanisms which may explain the phenotypic differences between patients and mice with *DNAJB11* or *PKD1* mutations. Our proteomics data sets in *Dnajb11*- and *Pkd1*-deficient cells provide novel candidate proteins such as COL4A1 or SDF2 for future studies investigating disease-relevant client proteins as well as additional co-factors in the chaperone network contributing to the maintenance of properly shaped kidney tubules.

In conclusion, our study shows that biallelic loss of *Dnajb11* in mice causes cystic kidney disease and fibrosis, mirroring the pathological manifestations observed in the corresponding human disorder. The absence of an overt phenotype in mice with monoallelic inactivation of *Dnajb11* supports a cellular recessive mechanism of cyst formation in this atypical form of ADPKD. Our findings demonstrate that proximal tubules are the predominant origin of cysts in *Dnajb11*-deficient mice, with postnatal conditional inactivation showing a marked reduction in disease severity compared to constitutive inactivation of *Dnajb11*. Furthermore, our findings indicate that loss of DNAJB11 impairs GPS cleavage of PC1 providing a plausible mechanism of cystogenesis. These insights contribute to a better understanding of the molecular mechanisms underlying cystic kidney disease and may help to develop therapeutic approaches aiming at modulating chaperone function to improve folding of mutant PC1. The potential benefit of this strategy in ADPKD has recently been demonstrated (50).

## Materials and methods

### Mouse strains and procedures

All experiments were conducted under permission G-19/149 of the Regierungspräsidium Freiburg, Germany. Animals were kept under specific pathogen-free conditions in the animal facility of the University Medical Center Freiburg according to institutional guidelines at 21-23° C, 45–60% humidity and a 12 h dark/light cycle. Unless otherwise stated mice received standard diet and water *ad libitum.* Frozen sperm of the LacZ-expressing knockout-first mouse strain *Dnajb11^tm1a^* (C57BL/6N-Dnajb11<tm1a(EUCOMM)Wtsi>) was obtained from EUCOMM (Helmholtz Zentrum München, Germany [EM:09265]) and used for *in vitro*-fertilization in C57BL/6N female donors.

*Dnajb11^fl^* mice (*Dnajb11^tm1c^*): Breeding of *Dnajb11^tm1a^* and *Act-FLP* (Tg(CAG-Flpo)1Afst) mice resulted in conditional *Dnajb11^fl^* mice (*Dnajb11^tm1c^*) which were subsequently bred with female B6.C-Tg(Pgk1-cre)1Lni/CrsJ mice to generate the constitutive *Dnajb11^−^* (*Dnajb11^tm1d^*) genotype. *Dnajb11^+/−^* mice were crossed with *Ksp-Cre* mice (B6.Cg-Tg(Cdh16-cre)91lgr/J) to generate *Ksp-Cre; Dnajb11^+/−^ mice which were backcrossed with homozygous Dnajb11^fl/fl^* mice to produce Ksp-Cre; *Dnajb11^fl/−^* and Ksp-Cre; *Dnajb11^fl/+^* littermates. Inducible *Pax8rtTA;LC1-Cre; Dnajb11^fl/fl^* animals (referred to as *Pax8-Cre; Dnajb11^fl/fl^)* were obtained from breedings with mice harbouring the *Pax8rtTA and LC1-Cre* transgenes (Tg(Pax8-rtTA2S*M2)1Koes, Tg(tetO-cre)LC1Bjd). For induction mice received doxycycline hydrochloride via the drinking water (2 mg/ml with 5% sucrose, protected from light) from post-natal day 28 (P28) to P42.

Sample collection and analysis: Both male and female 90 days-old mice were used in this study. Spot urine was collected from conscious animals. Blood from anesthetized mice (ketamine 160 mg/kg and xylazine 8 mg/kg) was sampled by retrobulbar venous plexus puncture and serum was obtained by centrifugation at 6000 g. Mice were killed by cervical dislocation and kidneys were perfused with 10 ml PBS and 40 ml PFA (3,7%) at a constant flow of 13 ml/min. Kidneys were embedded in paraffin for immunoflourescence and Hematoxylin/Eosin stainings. Embryos were harvested from pregnant female mice at 17,5 days post fertilization. Kidneys were either snap frozen for protein analysis or dissected and fixed for 24 h at 4° C in methyl Carnoy’s solution (60% methanol, 30% chloroform, and 10% glacial acetic acid). For blood and urine analyses the following kits were used according to the manufacturer’s instructions: Blood urea nitrogen (BUN): LT-UR; Labor&Technik, Eberhard Lehmann GmbH. Urinary creatinine: LT-CR Labor&Technik, Eberhard Lehmann GmbH.

### Imaging and image analysis

5 μm paraffin sections were stained with haematoxylin and eosin according to standard protocols. 2 µm paraffin sections were used for AFOG staining (Acid Fuchsin Orange G) following a standardized multi-step coloring protocol performed at the Institute of Surgical Pathology, Freiburg. 3-4 sections per mouse were imaged on an Axioscan 7 slide scanner (Zeiss) using a Plan-Apochromat 10x/0.45 objective and ZEN 3 software (ZEISS). In ImageJ software images were converted to 8 bit-greyscale and kidney cortex area was manually annotated. Tubular and cyst lumen was identified by thresholding grey-values 230-255. Cystic or dilation index was calculated as percentage of visible lumen/cortex area. Each dot in the graph represents the average dilation index of one mouse. For the quantification of KW/BW in Fig. 1H and cystic index in Fig. 1I, we excluded one outlier with excessively high KW/BW ratio and cystic index from the analysis according to Grubb’s outlier test, thus underestimating the actual increase of mean KW/BW ratio and cystic index. Immunolabeling was performed on 5 μm paraffin sections. After heat induced antigen retrieval (20 min at 96° C) in Tris-EDTA pH 9 (10 mM Tris, 1 mM Na_2_EDTA) and blocking (1 h; 1% BSA, 5% horse serum) sections were incubated with primary antibodies for Megalin/LRP2 (1:200, Novus Biologicals NB110-96417), Calbindin D-28K (1:200, Novus Biologicals NBP2-50028), Uromodulin (1:400, Abcam ab207170), Aquaporin-2 (1:100, SCBT sc-515770), smooth muscle actin (1:400, Agilent M08051). Secondary antibodies with different species reactivity were raised in donkey and conjugated to Alexa Fluor dyes (488, 555, 647; 1:600, Jackson Immuno Research). IF-imaging was performed on a Cell Discoverer 7 equipped with a LSM900 module (Zeiss).

### Cell Culture

IMCD-3 cells were cultured with DMEM:F12 (Lonza, cat. No. BE12-719F), fetal bovine serum was added to a final concentration of 10% and penicicillin/streptomycin (Sigma, P0781) to a final concentration of 1%. Subculturing of cells was performed twice a week in a 1:10 ratio. Cells were maintained in a humidified incubator at 37° C and 5% CO_2_.

HEK 293T cells were cultured with DMEM (Lonza, cat. NO. BE12-640F) fetal bovine serum was added to a final concentration of 10% and penicillin/streptomycin (Sigma, P0781) to a final concentration of 1%. Subculturing of cells was performed three times a week in a 1:10 ratio. Cells were maintained in a humidified incubator at 37° C and 5% CO_2_.

### Genome Editing

Genome editing of IMCD3 cells was performed as described before(51). *Dnajb11* knockouts were generated from an isogenic monoclonal mIMCD3 WT cell line to reduce clonal variability (38). To investigate the role of *Dnajb11* in PC1 cleavage and biogenesis, the gene was also knocked out in IMCD3 cells in which an N-terminal 3xFlag-tag and a C-terminal 2xV5-tag had been added to the endogenous *Pkd1* locus(51). Two gRNAs targeting Exon 1 (targeting sequence GGGCGCGCTACCTGTCTCCG) and Exon 10 (targeting sequence GAACTGGACTTTGAAAGAGG) were cloned in BPK1520 (Addgene #65777) and transfected with Amaxa Nucleofector II together with BPK4410, (for expression of HypaCas, Addgene #101178). Screening for knockout clones was performed via PCR and Sanger sequencing, knockout clones were confirmed via Western Blot.

### Immunoblotting

Cells were harvested five days after epithelial confluency or 24 h after transfection (HEK 293T). Samples were lysed in cold lysis buffer (1% Triton X-100, 20 mM Tris-HCl (pH 7.5), 50 mM NaCl, 50 mM NaF, 15 mM Na_4_P_2_O_7_, and 0.1 mM EDTA (pH 8)) supplemented with 2 mM Na_3_VO_4_ and complete protease inhibitor mixture (Roche). Lysates were first centrifuged at 4 °C for 15 min at 20.000 g and subsequently subjected to ultracentrifugation at 4° C for 30 min at 100.000 g. Supernatants were denatured 30 min at 42° C (for PC1 detection) or 5 min at 95° C with 2x Laemmli. Protein samples were separated by SDS-PAGE using precast Mini-PROTEAN TGX 4-15% (BioRad) or 3-8% Tris-Acetate gels (Invitrogen) and transferred onto PVDF membranes using a wet blot system for 1 h at 100 V (Biorad). Membranes were blocked with BSA and incubated with primary antibody overnight at 4° C. After incubation with secondary antibody and washing 3× 10 min, chemiluminescence was detected by a 16-bit ChemoCam system (Intas). Primary antibodies used were: anti-DNAJB11 (St. Cruz, sc-271240), anti-Gamma tubulin (Sigma, T6557), anti-V5 (BioRad, MCA1360), anti-PC1 (St. Cruz, sc-130554), anti-Vinculin (Cell signalling, #4650), anti-Flag (Sigma, F4049). Primary antibodies were used at 1:1000. Secondary antibodies (anti-rabbit, GE Healthcare NA934; anti-mouse, Dako P0447) were used at 1:10,000.

### Densitometry and statistical analysis

Densitometry was performed with ImageJ 1.46R. For quantitative analysis, the intensity of the protein of interest band was normalized to the intensity of the corresponding control band. All results are expressed as mean ± SEM. A p-value < 0.05 was considered significant unless otherwise specified. Statistical analysis was performed as described in the figure legends. Software used for statistics was GraphPad Prism 10.

### Immunoprecipitation

Protein A-Sepharose beads (GE Healthcare) were incubated with 2 µg of antibody (anti-V5, Millipore, AB3792) in PBS for 90 min at 4° C. For anti-FLAG-IPs, preconjugated antibody-coupled beads (Sigma, A2220) were used. Beads were washed three times for 10 min at 4° C with PBS and then combined with the respective lysates. Immunoprecipitation took place for 2 h on a rotating wheel at 4 °C. After 3 washing steps (each 10 min, 4° C), proteins were eluted for 30 min at 42° C with 2x Laemmli sample buffer and 100 mM DTT.

### Plasmids

For rescue experiments human *DNAJB11* was cloned into pLXSN. Patient mutations were introduced using Q5 site-directed mutagenesis Kit (NEB). Transduction was performed as described previously (51).

### Mass spectrometry analysis

For proteome comparison, two independent IMCD *Dnajb11^−/−^* as well as two independent IMCD3 *Pkd1^−/−^* clones were investigated in comparison to the corresponding isogenic IMCD3 WT cell line. Respective cell lines were harvested 5 d after confluency. Cell pellets were resuspended in lysis buffer (5% SDS, 50 mM triethyl ammonium bicarbonate (TEAB; Sigma, T7408), pH 7.5). Afterwards samples were sonicated using a Bioruptor device (Diagenode, Liège, Belgium). Samples were centrifuged at 13000 g for 8 min and the supernatant used in the following steps. Proteins were reduced using 5 mM tris (2-carboxyethyl) phophine hydrochloride (TCEP) (Sigma; 75259) for 10 min at 95° C and alkylated using 10 mM 2-iodoacetamide (Sigma; I1149) for 20 min at room temperature in the dark. Following steps were performed using S-Trap micro filters (Protifi, Huntington, NY) following the manufacturer’s procedure. Briefly, first a final concentration of 1.2% phosphoric acid and then six volumes of binding buffer (90% methanol; 100 mM TEAB; pH 7.1) were added. After gentle mixing, the protein solution was loaded to an S-Trap filter and spun at 2000 rpm for 0.5–1 min. The filter was washed three times using 150 μL of binding buffer. Sequencing-grade trypsin (Promega, 1:25 enzyme:protein ratio) diluted in 20 µl digestion buffer (50 mM TEAB) were added into the filter and digested at 47° C for 1 h. To elute peptides, three step-wise buffers were applied: a) 40 μL 50 mM TEAB, b) 40 µl 0.2% formic acid in H_2_O, and c) 50% acetonitrile and 0.2% formic acid in H_2_O. The peptide solutions were combined and dried in a SpeedVac.

The peptide concentration was measured using BCA (Pierce, 23225) and 25 µg of each sample was transferred to a fresh microreaction tube. HEPES (pH 8.0) was added to a final concentration of 0.15 M. Samples were labeled using TMT-16-plex (Thermo Scientific) (52). After labeling all samples were combined and 100 µg were fractionated by high pH reversed phase chromatography (XBridge C18 column, 150 mm × 1 mm column containing 3.5 µm particles (Waters)). An increasing linear gradient of acetonitrile from 10 to 45% over 45 min at a flowrate of 42 µl/min was applied using an Agilent 1100 HPLC system. 36 fractions were collected and concatenated into 10 fractions, which were vacuum-concentrated until dryness and stored at − 80° C until LC–MS/MS analysis.

800 ng of peptides were analyzed on a Q-Exactive Plus mass spectrometer (Thermo Scientific, San Jose, CA) coupled to an EASY-nLCTM 1000 UHPLC system (Thermo Scientific). The analytical column was self-packed with silica beads coated with C18 (Reprosil Pur C18-AQ, d = 3 Â) (Dr. Maisch HPLC GmbH, Ammerbusch, Germany). For peptide separation, a linear gradient of increasing buffer B (0.1% formic acid in 80% acetonitrile, Fluka) was applied, ranging from 5 to 40% buffer B over the first 90 min and from 40 to 100% buffer B in the subsequent 30 min (120 min separating gradient length). Peptides were analyzed in data dependent acquisition mode (DDA). Survey scans were performed at 70,000 resolution, an AGC target of 3e6 and a maximum injection time of 50 ms followed by targeting the top 10 precursor ions for fragmentation scans at 17,500 resolution with 1.6 m/z isolation windows, an NCE of 30 and an dynamic exclusion time of 35 s. For all MS2 scans the intensity threshold was set to 1.3e5, the AGC to 1e4 and the maximum injection time to 80 ms.

Raw data were analyzed with MaxQuant (v 1.6.14.0) with the built-in Andromeda peptide search engine (53) allowing two missed cleavage sites, no variable modifications, carbamidomethylation of cysteines as fixed modification, PIF was set to 0.75, and 16 plex TMT as isobaric label. The Mouse-EBI-reference database was downloaded from https://www.ebi.ac.uk/ on Feb 4th 2020. Only unique peptides were used for quantification.

Data were normalized on peptide level by equalizing the medians across all the channels and MS runs using the MSstatsTMT package (v. 1.8.2) in R (v. 4.0.3). Subsequently, protein intensities were log_2_ transformed. To identify differentially expressed proteins, we used the limma package (v. 3.46.0) in R using the “robust” method. P-values were adjusted using the Benjamini-Hochberg procedure. GO term analysis was performed using ShinyGo v0.77 (54). Tables S2 – S4 contain all proteins that meet significance- and fold-change thresholds (for adj. p: log_10_ > 1.3; for fold-change: log_2_ < −0.58 or log_2_ > 0.58).

For PC1 interaction studies, HEK 293T cells were transfected with pcDNA3.FLAG-PC1 (55), untransfected cells were used as control. After immunoprecipitation using anti-Flag antibody coupled beads, mass spectrometric analysis and data analysis was performed as described before (56). Three independent experiments were performed. For Fig. S1 and Table S1, only proteins with at least one peptide count in each n of the Flag.PC1-transfected cells were included in the analysis.

## Authors contributions

T.B., B.N., L.S., I.C., L.K., O.W., F.S., S.T., A.H. and L.W. conducted experiments. T.B., B.N., S.T., A.H., C.S., O.S., A.K. and M.K. analyzed the data. T.B., B.N., A.K. and M.K. designed the study and wrote the manuscript. T.B. and B.N. share first authorship because of similar contributions to the study, first mention of T.B. is based on him initializing the project.

## Acknowledgements and funding sources

The authors acknowledge Simone Diederichsen, Celine Costa and Andreas Ungi for expert technical assistance and Mariam Semmo for support with the PC1 affinity purification. We would like to thank the Lighthouse Core Facility for assistance with microscopy and cell sorting. Lighthouse Core Facility is funded in part by the Medical Faculty, University of Freiburg (Project Numbers 2023/A2-Fol; 2021/B3-Fol, the DTK and the DFG (Project Number 450392965). The work of O.S., C.S., A.K., and M.K. was funded by German Research Foundation (DFG) project ID 431984000 (SFB 1453). M.K. was supported by German Research Foundation (DFG) project ID 239283807 (TRR 152) and Germany’s Excellence Strategy (CIBSS, EXC-2189, project ID 390939984). The Proteomic Platform – Core Facility was supported by the Medical Faculty of the University of Freiburg to Prof. Dr. Oliver Schilling (2021/A3-Sch and 2023/A3-Sch).

## Supplementary Figures

**Fig. S1.**
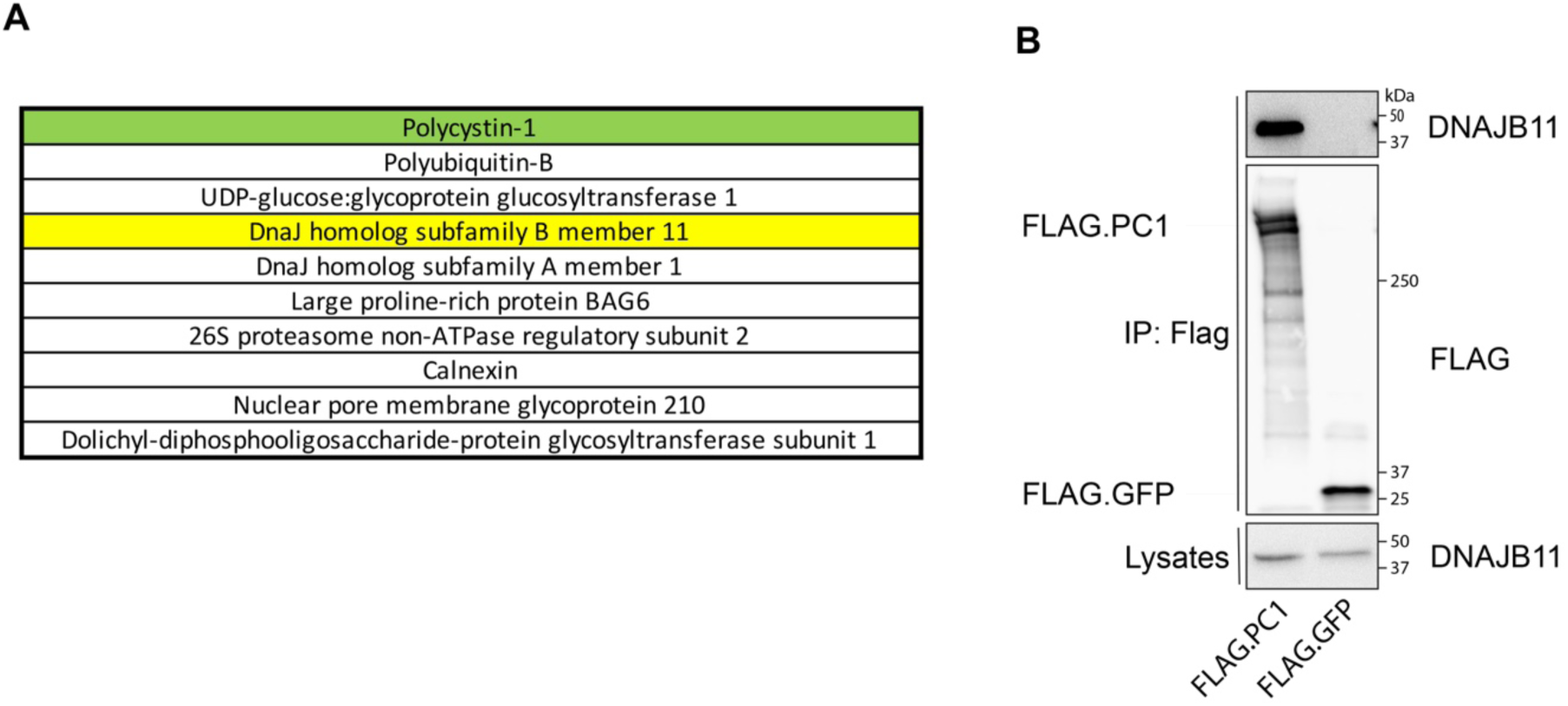
DNAJB11 interacts with PC1. **(A)** Affinity purification of FLAG-tagged PC1 followed by high-resolution mass spectrometry identifies co-purified proteins. List of 10 most enriched PC1 interacting proteins relative to untransfected control. Order is based on ratio and abundance of peptides (see Table 1). **(B)** In HEK 293T cells endogenous DNAJB11 interacts with overexpressed FLAG.PC1, but not with FLAG.GFP.

**Fig. S2.**
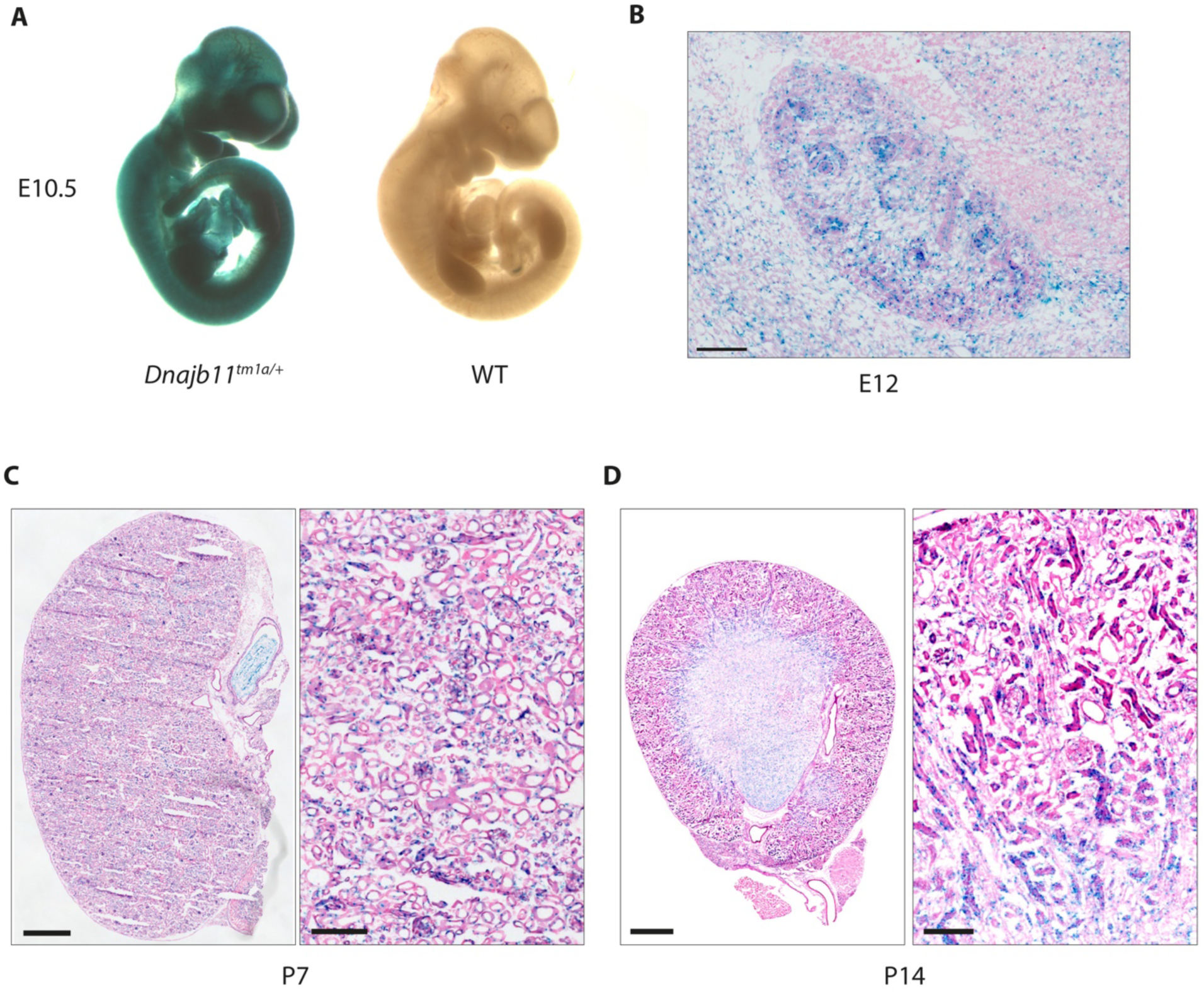
Detection of DNAJB11 expression via X-Gal staining of *Dnajb11^tm1a/+^* tissue samples. **(A)** X-Gal staining of whole embryo (E10.5). **(B-D)** X-Gal staining of kidney sections at indicated time points. Scale Bar: 100 µm (B), 500 µm (C left), 100 µm (C right), 500 µm (D left), 100 µm (D right).

**Fig. S3.**
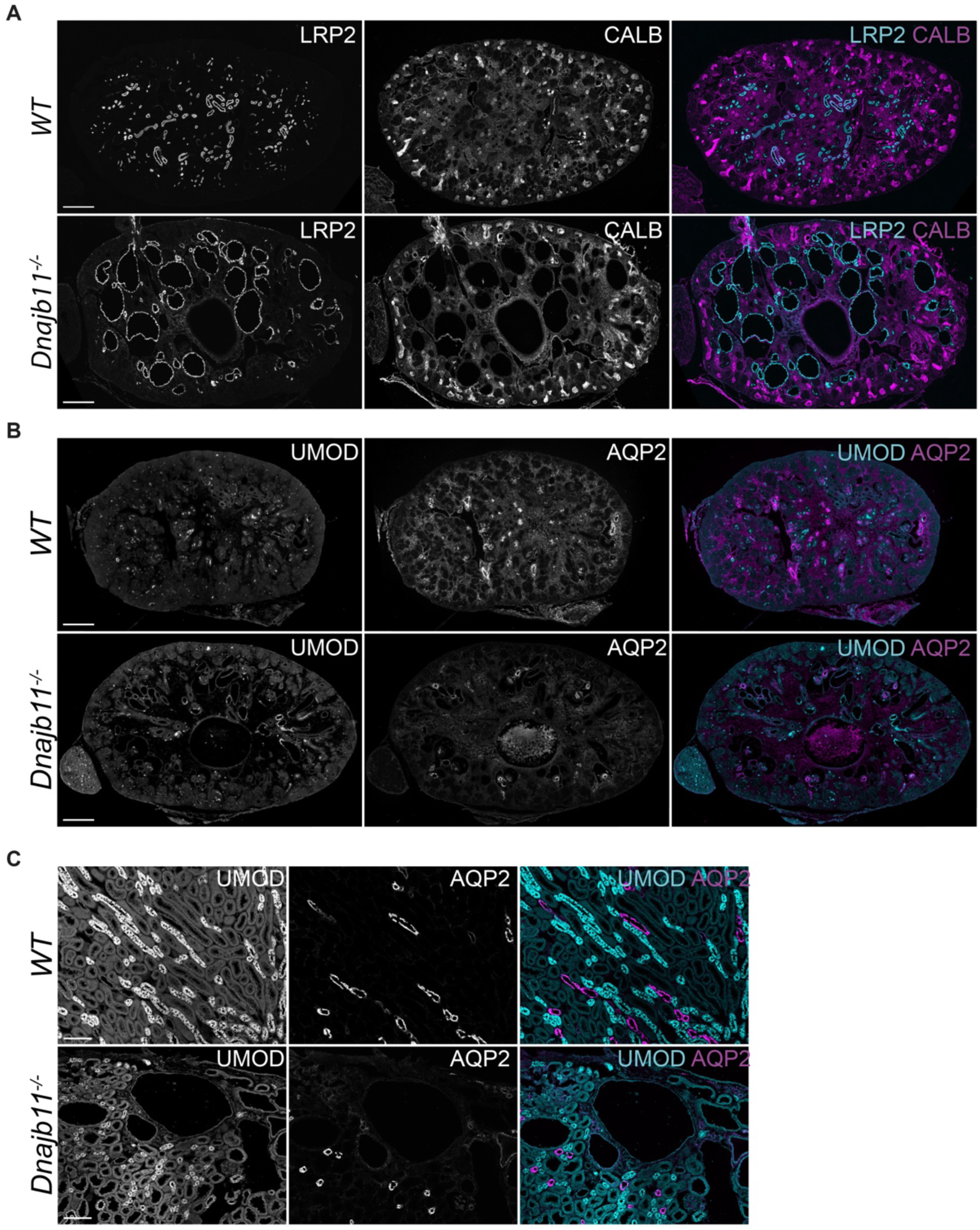
Immunohistochemistry using tubule segment markers in *Dnajb11^−/−^* kidneys. **(A), (B)** E17.5 WT and *Dnajb11^−/−^ k*idney sections were labeled with kidney tubule segment markers LRP2, CALB (A), UMOD and AQP2 (B). At E17.5, cyst-lining epithelium is exclusively labeled by LRP2. **C** P90 WT and *Dnajb11^−/−^ k*idney sections were labeled with kidney tubule segment markers UMOD and AQP2. Cyst-lining epithelium is not labeled by UMOD or AQP2. Scale Bar: 200 µm (A, B), 100 µm (C).

**Fig. S4.**
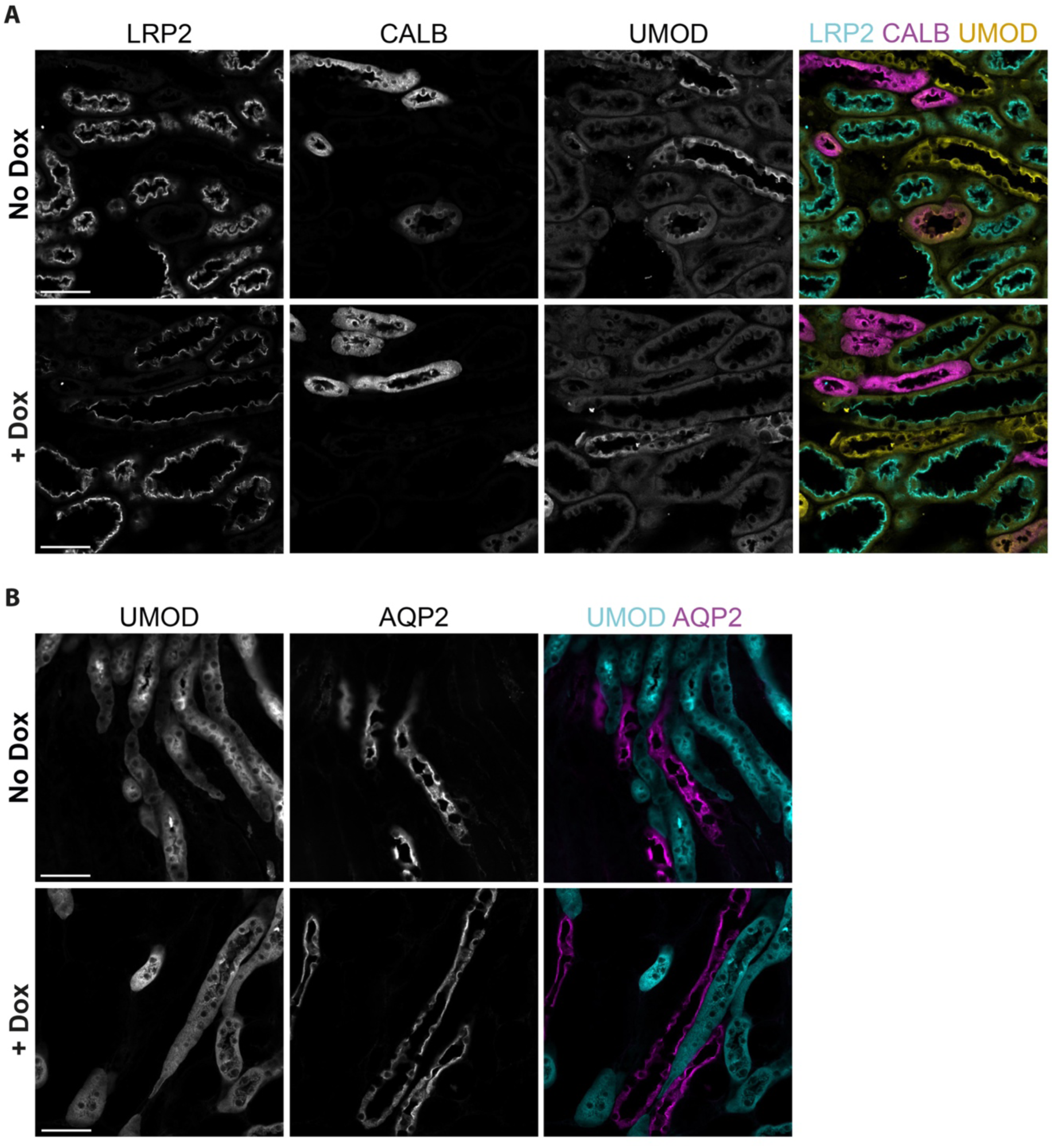
Immunohistochemistry using tubule segment markers in *Pax8; Dnajb11^fl/fl^* kidneys. **(A), (B)** Kidney sections of P126 doxycycline-induced *Pax8; Dnajb11^fl/fl^* as well as non-induced *Pax8; Dnajb11^fl/fl^* mice were labeled with kidney tubule segment markers LRP2, CALB, UMOD (A) and UMOD and AQP2 (B). Tubular dilations were observed exclusively in LRP2-positive tubular segments. Scale bar: 200 µm.

**Fig. S5.**
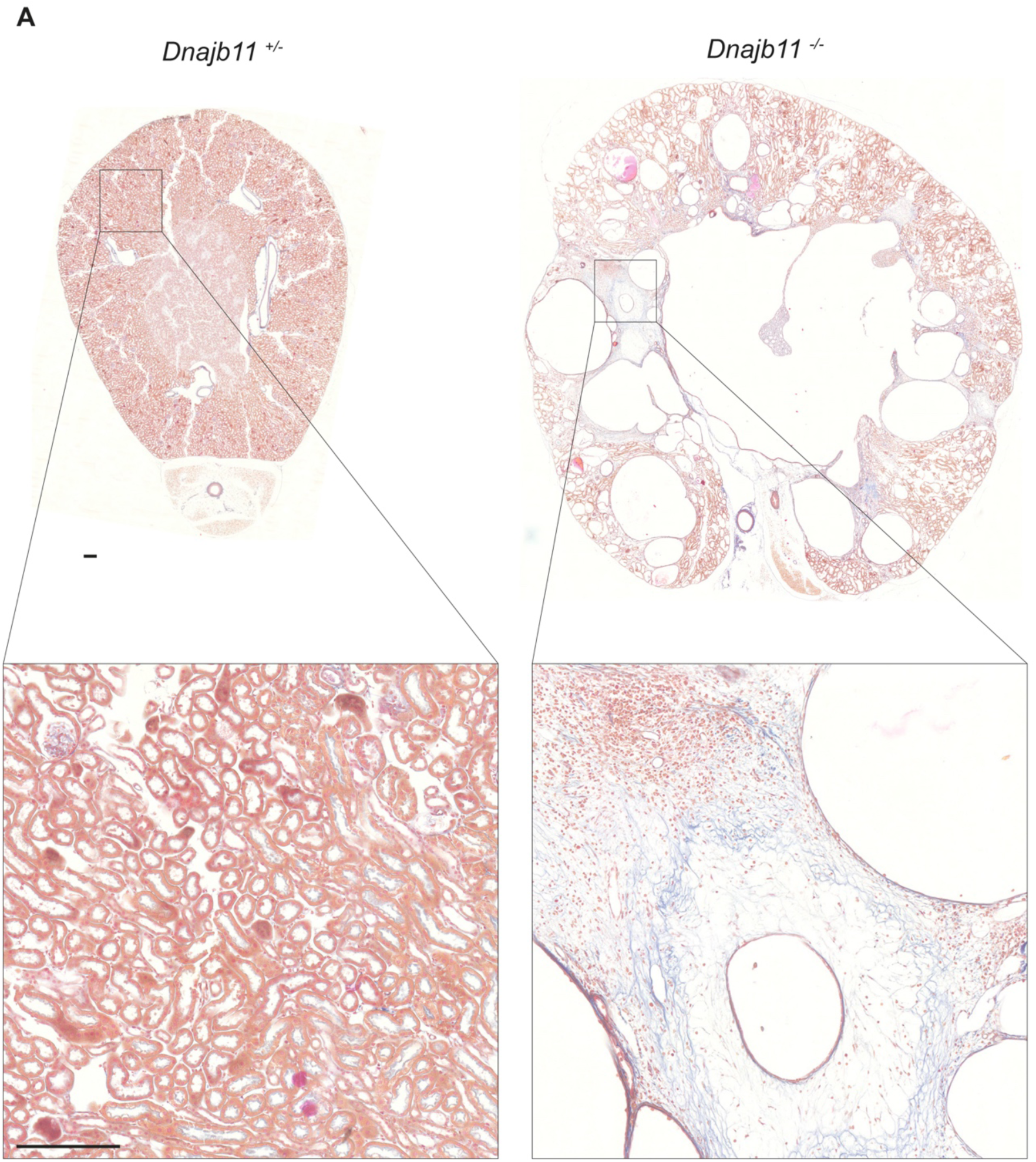
Acid Fuchsin staining of *Dnajb11^−/−^* kidneys. **(A)** Acid Fuchsin staining of kidney sections of P90 *Dnajb11^+/−^* and *Dnajb11^−/−^* mice shows pericystic fibrotic areas in *Dnajb11^−/−^* kidneys. Scale bars: 200 µm.

